# Spatial characterization of microbiota profiles along the gut in a widely used mouse model: effects of high-fat diet and fructooligosaccharides

**DOI:** 10.1101/2025.06.08.658095

**Authors:** Paul Taillandier, Lise Voland, Jean Debédat, Alix Archambeau, Sébastien Dussaud, Chloé Amouyal, Rafael Patiño-Navarrete, Federico Rey, Solia Adriouch, Eugeni Belda, Karine Clément, Tiphaine Le Roy

## Abstract

The intestinal microbiota plays a pivotal role in regulating metabolic processes, and its imbalance is linked to metabolic disorders. Modulating the gut microbiota composition and function through prebiotic supplementation has been repeatedly shown to improve host metabolism. While most studies have focused on the fecal microbiota due to the ease of sampling, fecal samples do not reflect the microbial dynamics throughout the gastrointestinal tract, as regional environmental conditions shape distinct microbiota composition.

Given the metabolic significance of the proximal intestine and the potential influence of microbiota on these processes, we characterized the microbiota composition along the gastrointestinal tract in a widely used model for studying host-microbiota interactions: C57Bl6J male mice fed a high-fat diet (HFD) with or without prebiotics supplementation (FOS, fructooligosaccharides).

The microbiota composition, determined by long-read Nanopore sequencing, was markedly altered by HFD and FOS supplementation not only in feces but also in the jejunum, ileum and caecum. In contrast to previous observations made in humans, obesity in mice was associated with a decrease in microbiome diversity in the small intestine, highlighting species-specific microbial responses to metabolic challenges. Additionally, the pronounced bifidogenic effect of FOS supplementation in the ileum suggests FOS fermentation in the small intestine in mice, contrary to what has been previously described in humans. Finally, we report a relative homogeneity of microbiome composition along the digestive tract, possibly due to the coprophagic behavior of mice. These findings challenge the translational relevance of rodent models for studying the role of the small intestinal microbiota in human metabolic disease.

## INTRODUCTION

The human body is composed of approximately equal proportions of human and microbial cells, the later comprising a diverse array of microorganisms, which belong to the bacteria, archaea, fungi, and protozoa kingdoms (1). In adults, the intestinal microbiota is estimated to comprise approximately 100 trillion bacteria, themselves belonging to an estimated number of approximately 4,600 species (2). These bacteria perform essential biological functions, including metabolization of xenobiotic compounds and non-digestible foods, the synthesis of several vitamins, the production of certain bioactive metabolites, and the maturation and activation of the immune system (3).

These key functions, including those yet to be described, enable it to exert a considerable impact on the host’s metabolism, as demonstrated and extensively studied in research conducted over the past 20 years. Indeed, an increasing body of evidence from both human and animal studies indicates a positive association between a diversified and balanced intestinal microbiota as well as metabolic and immune homeostasis (4,5). Conversely, a reduction in microbial diversity and an imbalance in microbial composition and functions lead to the disruption of essential metabolic processes and may promote low-grade inflammation. This alteration in the microbiota is now recognized as an important contributor to the pathophysiology of metabolic disorders (6).

Research exploring the bidirectional interplay between microbial ecology and the host’s metabolic profile is garnering increasing interest. Nevertheless, the majority of metagenomic studies primarily rely on fecal samples analysis. The microbial composition of these samples mostly reflects those of the distal colon, and differs significantly from the small intestinal microbiota in humans (7,8). These differences arise from the distinct physicochemical conditions in each segment of the digestive tract, including variations in pH, mucus composition, oxygen and nutrient availability, as well as factors such as bile acids secretion, and the production of antimicrobial peptides (9,10).

Microbial composition is not only shaped by regional conditions in the gut but also by dietary factors, particularly the amount of fermentable compounds in the intestinal lumen. Some of these compounds exert prebiotic activity, that is to say the enhancement of host health by selectively stimulating the growth and activity of beneficial bacteria with non-digestible compounds (11). Fructans exert beneficial effects on host metabolism by selectively stimulating the growth of beneficial bacteria of the gut microbiota such as *Faecalibacterium prausnitzii*, *Bifidobacterium*, *Lactobacillus* and *Akkermansia muciniphila*. For some of these species, their positive effects on host health are attributed to the production of short-chain fatty acids (SCFAs) (12). Among these prebiotics, fructooligosaccharides (FOS) are short-chain polymers comprising glucose and several fructose units. FOS supplementation in HFD-fed mice also strongly impact microbiome composition in the small intestine with a significant increase in *Bifidobacterium* genus abundance (13). Thus, given the alterations in microbiota composition within the small intestine, the beneficial effects of prebiotics may, at least in part, be mediated by mechanisms specific to this region of the digestive tract.

Studying the microbiota of the proximal small intestine is particularly important, as this region plays a crucial role in the detection, digestion and absorption of nutrients. Moreover, it plays a key role in activating metabolic pathways that influence energy expenditure, thermogenesis, satiety, food intake, and glucose homeostasis (14,15). Alterations in the function of the small intestine have been reported in obesity and associated metabolic disorders and are believed to contribute to the progression of these conditions as reviewed in (16). These alterations of the small intestine functions can be mediated by diet, a main determinants of intestinal homeostasis (17). For example, research has demonstrated that a western diet disrupts the mucus layer in mice, affecting not only the colon but also the small intestine (18). In addition, HFD has been shown to inhibit the intestinal PPAR-γ pathway in mice, leading to a disruption in the microbial ecosystem of the small intestine, as well as impairing its barrier function (19). Furthermore, HFD alters small intestinal microbiota composition and promotes the absorption of triglycerides and cholesterol in mice (20). These effects are possibly mediated by specific microbial products, since it has been demonstrated *in vitro* that the supernatant of certain intestinal bacteria can modulate lipid metabolism in enterocytes. (21,22).

Mouse models of diet-induced obesity and prebiotics supplementation are very popular models to study the role of the microbiota in metabolic diseases. A few studies, primarily based on 16S rRNA gene sequencing, have described variations in the relative abundance of specific genera and broad shifts in beta-diversity in the mouse small intestine. Nonetheless, a high-resolution, region-specific taxonomic characterization of the intestinal microbiota, especially under HFD and prebiotic intervention, remains lacking despite the widespread use of this model to study interactions between the diet, the host and its intestinal microbiome. Given the major role of the small intestine in various metabolic processes and the potential influence of gut microbes, we sought to refine our understanding of a widely used mice model for host-microbiota interaction in metabolic diseases by providing an in-depth analysis and characterization of the microbiota composition along the digestive tract upon HFD and prebiotics supplementation.

To address this gap, mice were fed either a chow diet or a HFD, with or without FOS supplementation for 15 weeks. Jejunal, ileal and cecal contents as well as feces were collected from each experimental group, and microbiota composition at the species level was analyzed using long-read nanopore sequencing, enabling a comprehensive characterization of microbial shifts across intestinal regions.

## RESULTS

### FOS limits body weight gain and metabolic alterations induced by the HFD

As expected, mice fed a HFD gained significantly more body weight than those fed a chow diet (**Fig. 1a**). Body composition analysis by nuclear magnetic resonance showed that the increase in body weight was primarily due to an increase in fat mass rather than lean mass (**Fig. 1b and 1c**). As previously described, FOS limited the body weight and fat mass gain induced by the HFD (**Fig. 1a to 1c**) (23).

**Figure 1.**
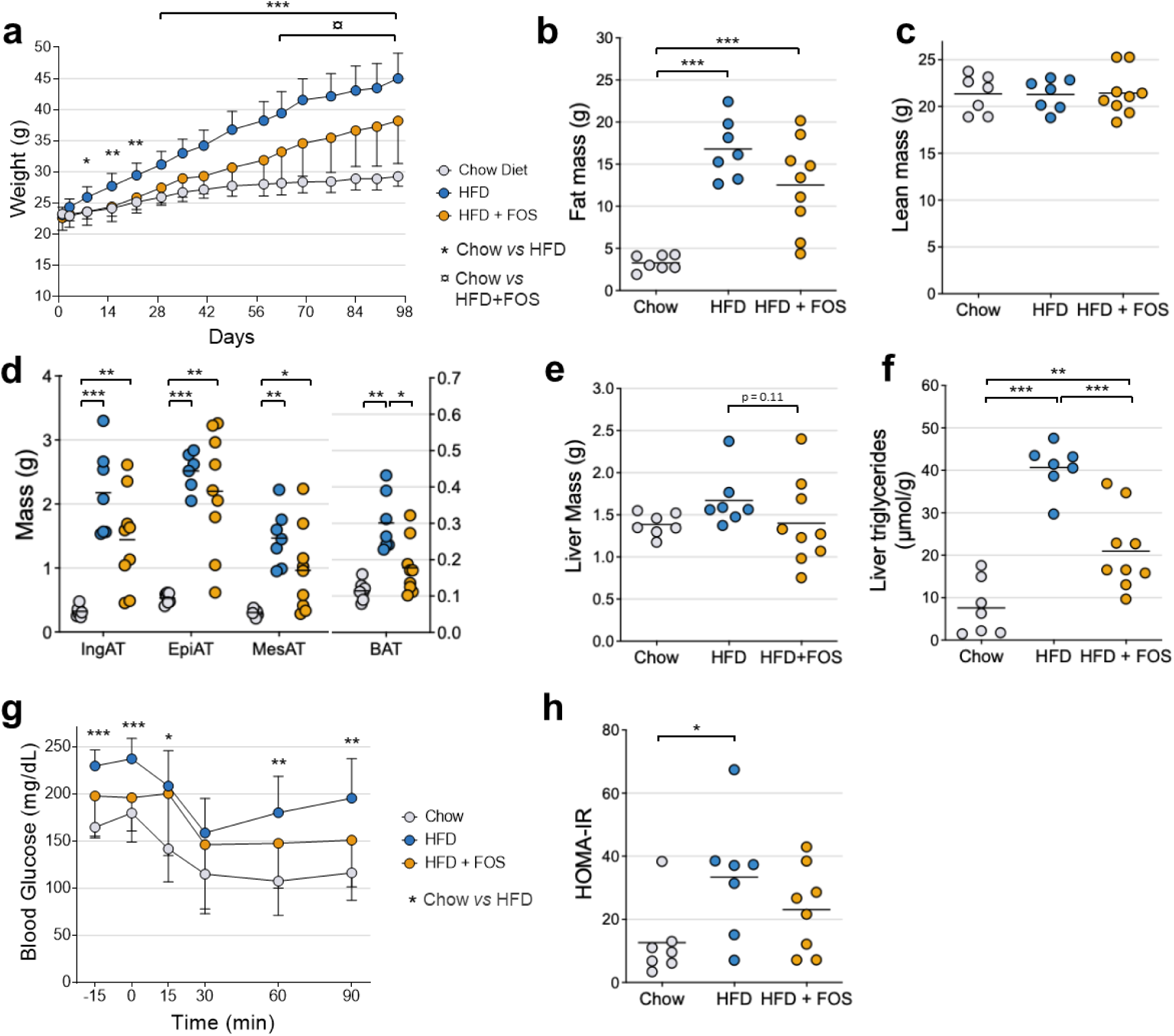
<u>Characterization of the effects of high-fat feeding and fructooligosaccharides on mice metabolism.</u> (**a**). Body weight of mice fed a chow diet or a HFD and supplemented with fructooligosaccharides (FOS) in drinking water (10% weight/vol). (**b**) Total fat mass, (**c**) lean mass, (**d**) adipose tissues and (**e**) liver weights of chow diet and HFD-fed mice treated with FOS for 13 weeks. (**f**) Liver triglycerides concentration of chow and HFD-fed mice treated during 13 weeks with FOS. (**g**) Insulin sensitivity test of mice after 8 weeks of chow diet or HFD and FOS supplementation. (**h**) Insulin resistance index (HOMA-IR) of mice after 12 weeks of chow diet or HFD and FOS supplementation. Number of mice per group: 7–9. Results are represented as dot plots with mean ± SD. Data for panels A and G were analyzed using a two-way ANOVA, while data for panels B-F and H were analyzed using a one-way ANOVA. All ANOVAs were followed by Tukey’s pairwise multiple comparison test. *: p ≤0.05; **: p ≤ 0.01; ***: p ≤ 0.001 for Chow diet vs HFD comparisons. In panel A, ¤: p ≤ 0.05 for Chow diet vs HFD + FOS comparisons.

Body composition measurements determined by Nuclear Magnetic Resonance (NMR) was corroborated by the measure of the weight of multiple fat pads, showing that mice on the HFD had significantly greater inguinal, epididymal, mesenteric, and brown adipose tissue (BAT) mass compared to controls (**Fig. 1d**). Fat pads weight was lower in mice supplemented with FOS although this effect was not statistically significant except for the interscapular brown adipose tissue. In HFD-fed mice, FOS treatment decreased liver weight (**Fig. 1e**) and HFD resulted in a 5.8-fold increase in liver triglycerides levels, which was counteracted by FOS supplementation (**Fig. 1f**).

The lower adiposity in mice fed the HFD and treated with FOS was associated with a non-statistically significant decrease in fasting blood glucose and improved insulin sensitivity, as shown by the insulin sensitivity test performed at week 8. (**Fig. 1g**). We also confirmed that FOS tend to limit the development of insulin resistance induced by the HFD, as shown by the calculation of the insulin resistance surrogate, HOMA-IR index (**Fig. 1h**). However, the effect of FOS on this index was not statistically significant.

Collectively, these results confirm the overall beneficial impact effect of FOS supplementation on the metabolic phenotype of mice in a model of HFD-induced obesity.

### Effect of diet and prebiotic treatment on the diversity of bacterial communities throughout the digestive tract and in the feces of mice

The microbiota composition along the digestive tract of the mice was determined using long-read Nanopore sequencing as previously described (24). We observed large discrepancies in the number of microbial reads between ecological niches **(Fig. S1a)**, leading to variations in sequencing depth. This was highlighted by statistically different slopes at the end of the rarefaction curves (**Fig. S1b**), indicating an uneven taxonomic coverage. Consequently, α-diversity analyses will be conducted on a “niche-by-niche” basis, rather than across the entire dataset, to avoid biases arising from uneven taxonomic coverage.

Several diversity indices reflecting different aspects of community structure: Richness (the number of observed species), Evenness (the uniformity of species abundances), as well as the Simpson and Shannon indices, which account for both richness and evenness were computed to assess the alpha diversity of bacterial communities in each sample. Compared to the Chow diet, the HFD feeding resulted in a decrease in alpha diversity indices throughout the digestive tract **(Fig. 2a to 2o, Fig. S2 and table 1**). Notably, the Richness index decreased more markedly in the jejunum (fold-change = 0.36) and ileum (fold-change = 0.33), than in the caecum (fold-change = 0.97) and feces (fold-change = 0.86). Prebiotic treatment partially restored bacterial diversity in the jejunum and ileum of mice fed a HFD, with significant effect observed in the ileum. Specifically, the restoration of the microbiota richness in the jejunum and ileum was 17% and 32%, respectively. Surprisingly, species richness was lower in the cecal content (p ≤ 0.05) and feces (p = 0.065) of FOS-treated HFD mice compared to untreated mice on the same diet. No restoration, even partial, of the Evenness and Shannon indices was observed in the caecum and feces of the mice after 13 weeks of FOS supplementation. Regarding the Simpson index, a non-significant decrease was observed in caecum and feces induced by FOS, suggesting a partial restoration of diversity according to this index.

**Figure 2.**
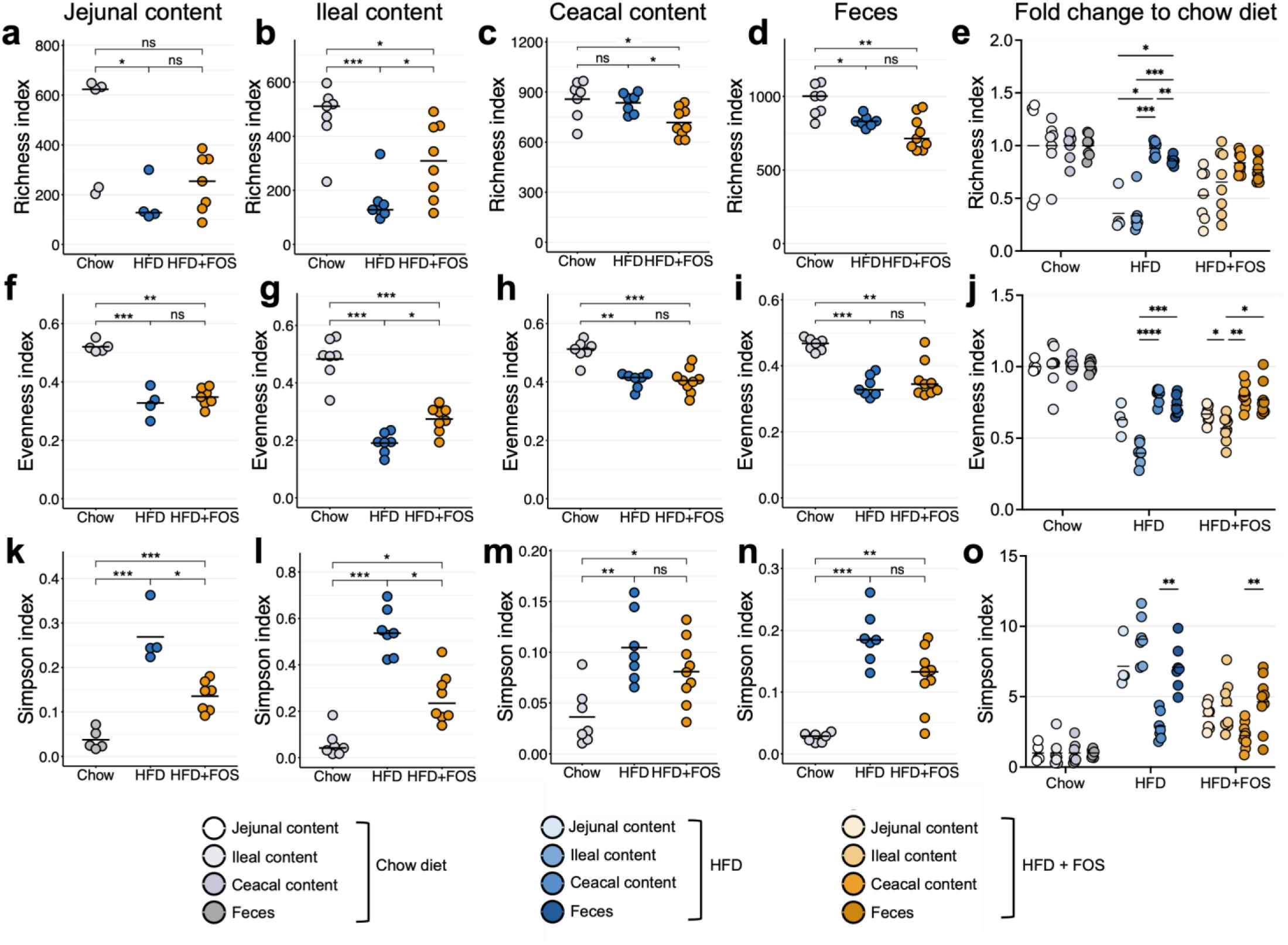
<u>High-fat diet and fructooligosaccharides supplementation modulate the alpha-diversity of the microbiome along the digestive tract.</u> (**a-d**) Richness, (**f-i**) Evenness and (**k-n**) Simpson index of the jejunal content, ileal content, cecal content and feces of mice fed a chow diet or a HFD and supplemented with fructooligosaccharides (FOS) in drinking water (10% weight/vol) for 13 weeks. The fold change was defined as the ratio between the measured value for (**e**) Richness (**j**) Evenness and (**o**) Simpson’s index in the HFD or HFD.FOS groups and the mean of the values of the same index observed in the chow diet group. Number of samples per group: 4–9. Results for c, f, g, k, m, e, j and o are represented as dot plots with mean, while the results for the other panels are represented as dot plots with median. Data for c, f, g, k, and m were analyzed using ANOVA followed by Tukey’s post-hoc test. For panels e, j, and o, Tukey’s multiple comparisons test was applied after adjusting for a mixed model. Data from the other panels were analyzed using the Kruskal-Wallis test followed by Dunn’s pairwise multiple comparison procedure. ns: p > 0.05, *: p ≤ 0.05; **: p ≤ 0.01; ***: p ≤ 0.001.

### Effect of a high-fat diet, FOS supplementation, and ecological niche on bacterial family composition

We examined the influence of HFD and FOS supplementation as well as gut segment on the relative abundance of the most abundant bacterial families among the five most dominant phyla. Statistical significance was assessed using a mixed-effects linear model in order to consider the non-independence of observations between the different ecological niches of the intestinal tract of each mouse. We observed a significant impact of diet, FOS supplementation and gut localization on the abundance of all 14 selected families (**Fig. 3 and S3, Table 2**).

**Figure 3.**
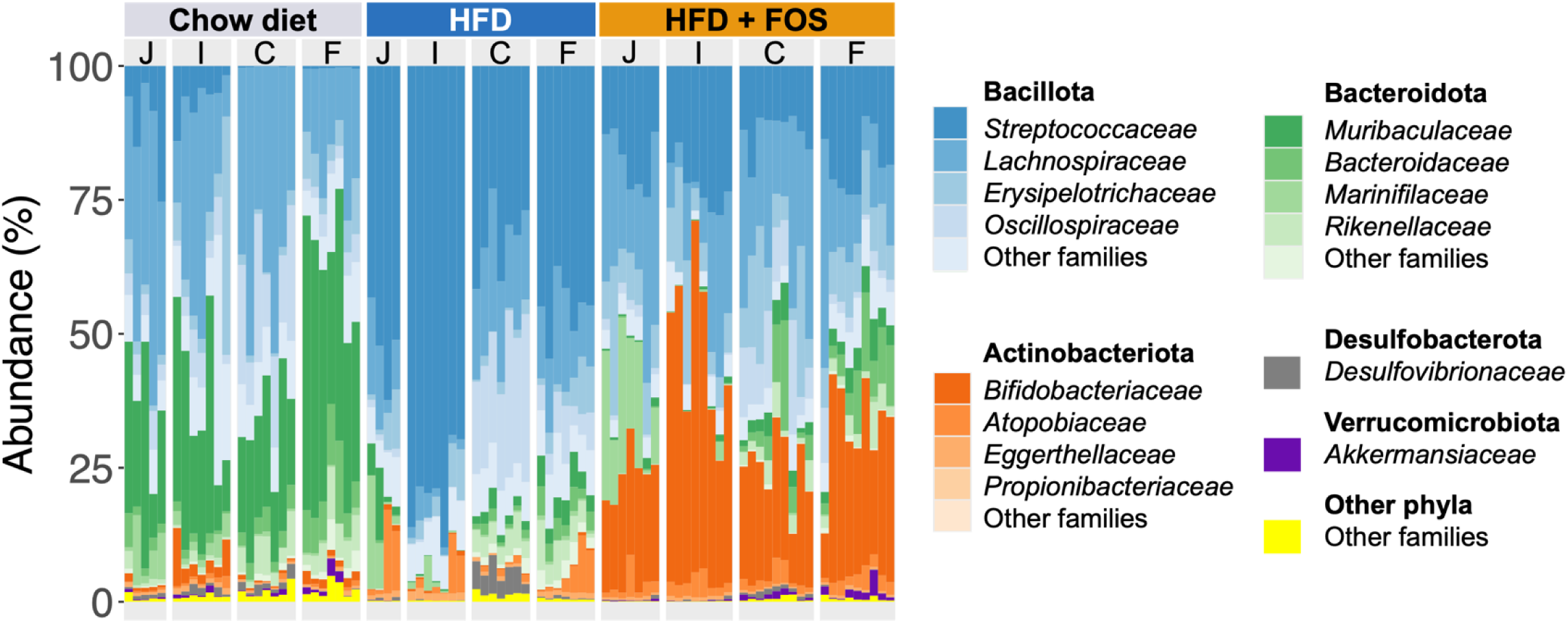
<u>Profile of dominant bacterial families among the five most dominant phyla along the digestive tract and in the feces of mice fed a control or HFD diet and supplemented with FOS:</u> Relative abundance (%, y-axis) of the most abundant bacterial families among the five most dominant phyla by group and ecological niche. Phyla were identified and color-coded, while bacterial families were represented by distinct color shades within each phylum. Each column represents a sample. Number of samples per group: 4–9. The letters J, I, C, and F, shown at the top of the graph within the facets, correspond to jejunal, ileal, cecal, and fecal samples, respectively.

Specifically, we found that the Bacillota phylum, especially the *Streptococcaceae* family, was significantly more abundant in the HFD group than in the Chow and HFD + FOS groups throughout the digestive tract. In contrast, the *Muribaculaceae* family (phylum Bacteroidota) showed a higher relative abundance in the Chow group compared to both the HFD and HFD + FOS groups.

We also observed notable changes in the relative abundance of 9 of the 14 dominant bacterial families depending on the ecological niches in mice fed the same diet (**Fig. 3 and S3, Table 2**). For example, in HFD-fed mice supplemented with fructooligosaccharides, the abundance of *Marinifilaceae* was significantly higher in the jejunum, than in the ileum, caecum, and feces (Tukey’s post-hoc test). In contrast, the *Rikenellaceae,* the *Muribaculaceae* and the *Bacteroidaceae* families were more predominant in the distal intestine (caecum and feces) than in the jejunum and ileum of the mice across all three experimental groups. Furthermore, the *Desulfovibrionaceae* and *Oscillospiraceae* families were primarily present in the caecum of mice from all three groups. Notably, *Bifidobacteriaceae* family (phylum Actinobacteriota) was less abundant in the HFD group compared to the control diet group but showed a marked increase with FOS supplementation, particularly in the ileum. *Akkermansiaceae* family was predominantly found in the distal gut of Chow-fed mice. Its abundance was slightly, although not significantly, decreased by the HFD but was partially restored by FOS treatment.

In summary, both the diet and ecological niche play significant roles in shaping the relative abundance of these bacterial families.

### Diet and ecological niche influence the composition of the microbiota throughout the digestive tract

To assess the impact of diet and ecological niche on the ecosystem structure of the microbiota, we performed beta diversity analyses based on the Bray-Curtis dissimilarity matrix and visualized samples similarity through non-metric multidimensional scaling (nMDS) in each ecological niche. In each of these ecological niches, we observed a relatively high level of intra-group similarity among the samples (**Fig. 4a to 4d**). In contrast, the dissimilarity is significantly higher between samples from mice fed with different diets within each ecological niche. The visualization of all the samples together highlights that samples primarily cluster together based on the diet fed to the mice, forming three distinct groups, rather than on the ecological niche (**Fig. 4e**). The samples from mice fed the same diet but from different ecological niches exhibit strong similarities. These findings suggest that diet and prebiotic supplementation have a more significant impact on the gut microbiota composition than the ecological niche. To confirm this visual observation, we compared the average Bray-Curtis distances between pairs of samples from different ecosystems within the same mice group, with those calculated between pairs of samples from different experimental groups within the same ecological niche (**Fig. 4f** and **Fig. S4**). Our analysis shows that the Bray-Curtis dissimilarity due to diet and FOS supplementation (average 0.73) is significantly higher than that attributed to the ecological niche (average 0.51). These results quantitatively confirm that diet and prebiotics supplementation have a significantly stronger impact on gut microbiota composition than the sampling location within the gastrointestinal tract in mice.

**Figure 4.**
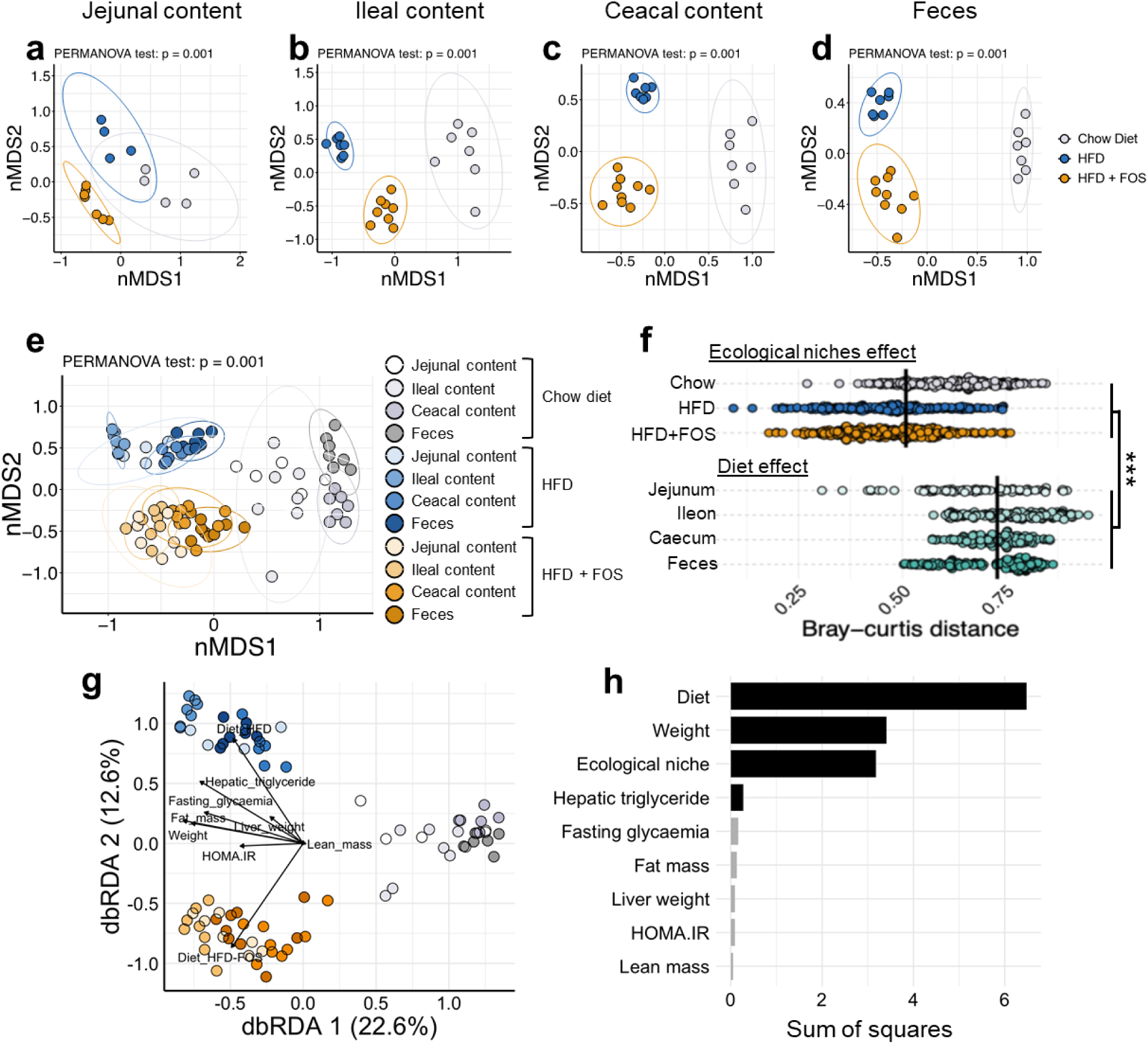
<u>High-fat diet, fructooligosaccharides supplementation and digestive tract localization significantly modulate microbiome beta-diversity</u>. Animals were fed chow or HFD and supplemented with fructooligosaccharides in drinking water (10% wt/vol) for 13 weeks. Non-metric Multi-Dimensional Scaling (nMDS) plots based on the Bray-Curtis distance of the microbiome in (**a**) jejunal content, (**b**) ileal content, (**c**) cecal content, (**d**) feces, (**e**) all samples. (**f**) Dot plot showing Bray-Curtis dissimilarity distances between pairs of samples from different ecosystems but within the same group and those from different experimental groups but the same ecological niche. Student test. ***: p ≤ 0.001. (**g**) Distance-based redundancy analysis (dbRDA) plots based on the Bray-Curtis distance of all samples. Arrows illustrate both the direction and magnitude of the explanatory power of each variable on the microbiome variability. (**h**) Variables correlating the most to microbiome compositional variation independently (univariate effect size). Significant variables (p ≤ 0.05) are indicated by black bars, while non-significant variables are represented by gray bars. Number of samples per group: 4-9.

We next performed a distance-based redundancy analysis (dbRDA) **(Fig. 4g)** in order to evaluate the explanatory power of phenotypic variables on variations of the microbiome composition. The variables “Diet”, “Fasting glycaemia”, “Body weight”, “Fat mass”, “Hepatic triglycerides”, and “Liver weight”, were identified as having the largest explanatory power, represented by long arrows pointing towards the samples from HFD fed mice. This indicates a significant association of these metabolic parameters with the microbiome structure of the mice along the digestive tract. In contrast, lean mass showed minimal influence on sample’s diversity. For further investigation, an ANOVA was performed on the dbRDA results to statistically assess the relative contribution of each explanatory variables to the variations of microbiome composition **(Fig. 4h)**. Consistent with previously observed results, diet significantly explained the variance in microbiome composition, with a more pronounced effect size than that attributable to biological niche in the digestive tract. Unlike other metabolic variables, both weight and hepatic triglyceride levels were significant, with weight showing the most pronounced effect size after diet.

### Minimal ecological niche effect on the gut microbiota composition may be driven by coprophagy

Although the ecological niche significantly influences the overall microbiota structure, we initially hypothesized that the distinct environmental conditions in each digestive tract segment would lead to greater taxonomic differences between these segments and that the effect of the segment would outweigh the effect of the diet. The relatively low dissimilarity observed between gut segments suggests that microbial composition may be homogenized, likely due to the coprophagic behavior of mice. This behavior could facilitate the transfer of microbial species from feces to the proximal intestine, reducing regional differences. To further explore this hypothesis, we computed Venn diagrams to determine species overlap within each diet across ecological niches. A species was considered present in an ecological niche within a group of mice if detected in at least 20% of the samples. Then, we identified the species shared across the three diets for each ecological niche and visualized their distribution using Venn diagrams (**Fig. 5a to 5d and table 3**), in order to determine the number of specific and shared species between different gut segments and feces.

**Figure 5.**
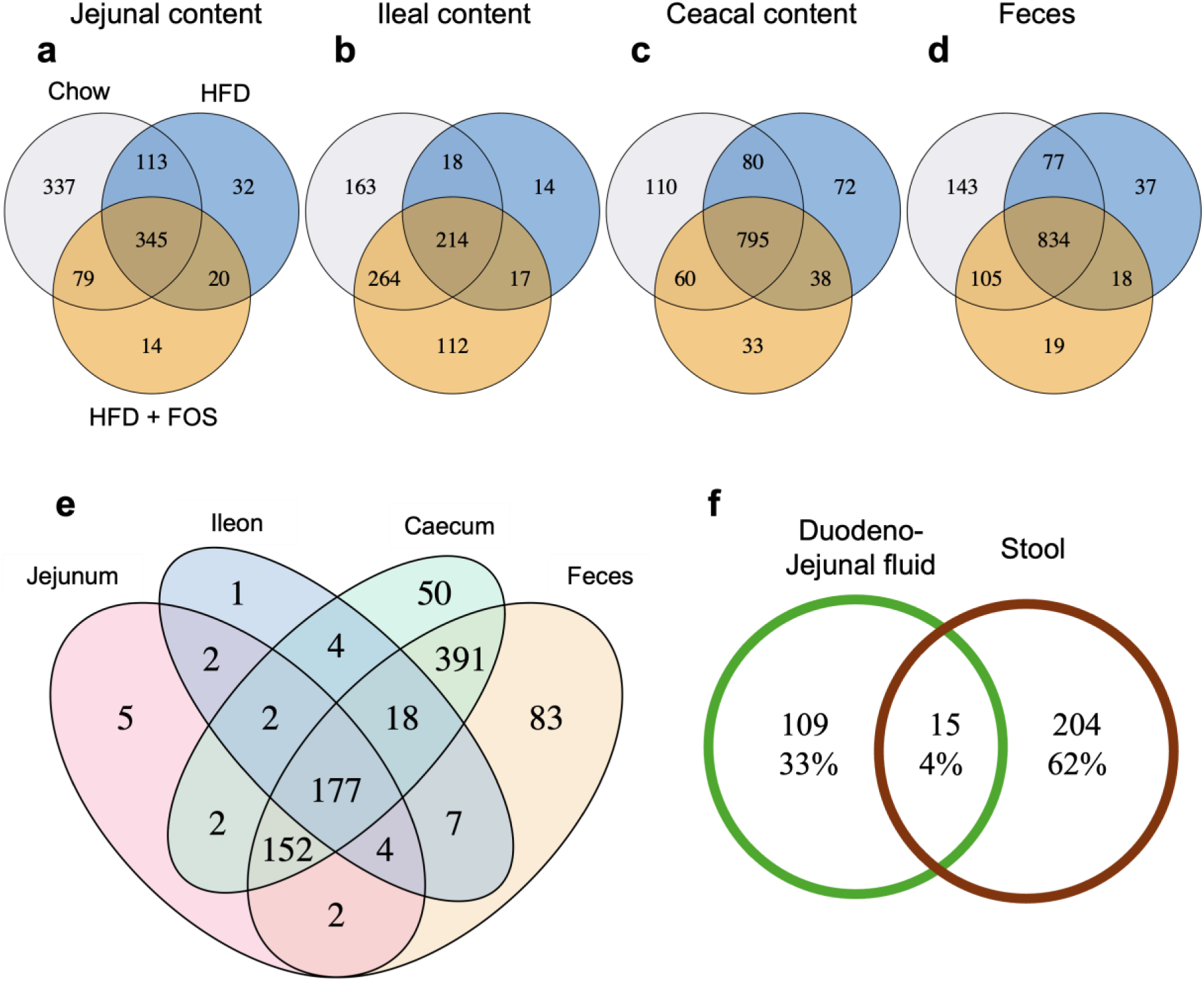
<u>Venn diagrams showing the unique and shared species</u> within each dietary regimen for (**a**) jejunal, (**b**) ileal, (**c**) cecal and (**d**) feces. A species was considered present if it was detected in at least 20% of the samples within a given group. (**e**) Venn diagram showing the unique and shared species between the four ecological niches within the 900 species common to the three diets groups. (**f**) Venn diagram illustrating the unique and shared bacterial species between duodenojejunal fluid (green) and stool (brown) in 30 human participants (15 obese and 15 normal weight). Duodenojejunal fluid was collected via endoscopy, and microbial composition was analyzed using Illumina sequencing and MetaPhlan 4 pipeline. A prevalence threshold of 20% was also applied for species inclusion. Results are presented as both absolute counts and relative abundance (% of total species).

Our analysis showed that the jejunal and ileal regions shared fewer taxa between the three diets groups than the fecal and cecal regions. Specifically, 345 (36%) species were shared across the three diets in the jejunum, 214 (26%) in the ileum, while 795 (67%) species were shared in the caecum and 834 (68%) in the feces (see **Table S1** for the list of species in each section of the Venn diagrams). The caecum and feces regions had the lowest number of exclusive species for the Chow diet group, with only 110 and 143 species respectively, compared to 337 in the jejunum and 163 in the ileum. The number of species exclusive to the HFD+FOS diet was higher in the ileum (113 species) than in the jejunum (14), caecum (33) and feces (19). These findings suggest that dietary effects on taxon presence/absence are more pronounced in the small intestine than in the more distal parts of the digestive tract in mice.

We then selected the taxa shared across the three diets for each ecological niche and computed a Venn diagram based on the four ecological niches (**Fig. 5e**). A total of 900 taxa were included in this analysis, of which few species were detected exclusively in the jejunal and ileal regions, with only 5 and 1 species, respectively. In contrast, the cecal and fecal regions hosted 50 and 83 unique species, respectively. A total of 152 species (17% of the total) were shared between the jejunum, caecum, and feces while only 18 species were common to all three distal regions (ileum, caecum, and feces). The largest overlap occurred between the caecum and feces, with 391 shared species, representing 43% of the total species selected. Finally, 177 species were common to all ecological niches, and 335 species (37%) were shared either across all ecological niches or between the jejunum and at least one distal region (caecum and/or feces).

This large overlap of bacterial species present in the proximal and distal intestinal regions of the gut may be attributed to the coprophagic behavior of mice. To further explore this hypothesis, we determined the number of shared taxa between the proximal small intestine and stools in human in order to evaluate the compositional similarity of the proximal and distal microbiome in the absence of coprophagy. In a previous study, we characterized the duodenojejunal and fecal microbiota of thirty individuals, of which half were affected by obesity (8). The analysis revealed a low bacterial species overlap between the proximal small intestine (duodenojejunal fluid) and stools in human. Specifically, only 4% (15 out of 328 species) in humans, which is significantly lower than the jejunal and fecal overlap in mice (37%, 335 out of 900 species) (**Fig. 5f**).

## DISCUSSION

In this study, we comprehensively investigated the modifications of the intestinal microbiota composition along the digestive tract induced by high fat diet and fructooligosaccharides supplementation. Our results show that in mice, the dynamics of the microbiota in the small and large intestine are highly similar, in contrast to what is observed in humans.

The small intestine is the main site of enzymatic digestion and nutrient absorption and its surface area is significantly larger than that of the large intestine (25), making it a crucial interface for host-bacterial interactions. Alterations in the microbiota composition of the small intestine have been associated with obesity phenotypes (8), coeliac disease (26), and stunting in children (27). Despite its importance in human health and the significant influence of the microorganisms that reside there, the microbiota in this region of the digestive tract remains understudied and poorly described, due to the challenges of accessibility to samples.

Mice fed a HFD and supplemented with FOS are widely used to study host-microbiome interactions in metabolic diseases (28–31). Prebiotics primarily exert their beneficial effects by modulating gut microbial ecology and promoting the production of bioactive metabolites such as short-chain fatty acids. However, most studies focus on fecal microbiota, while the composition and function of the small intestinal microbiota remain largely unexplored. Although non-digestible fibres exhibiting prebiotic activity are generally assumed to be fermented in the distal parts of the gut, the effects of prebiotics in the small intestine are poorly characterized. Herein, we characterized microbiota composition across the jejunum, ileum, cecum, and feces using long read nanopore sequencing. Sequencing depth in the proximal small intestine was significantly lower than in the distal segments of the gut, probably due to the naturally lower microbial load in this segment of the gut. This constitutes a limitation of the present study as we were consequently not able to perform a rigorous comparison of the alpha-diversity of the microbiome in the different segments of the gut. Future studies could potentially improve sequencing depth through additional sequencing effort or the use of alternative animal models such as rats to obtain greater sample amounts.

We confirmed that prebiotic supplementation mitigates HFD-induced fat mass gain and alterations in lipid and carbohydrates metabolism (32,33). High fat diet significantly reduced the microbiome alpha diversity, with a more pronounced effect in the jejunum and ileum compared to the caecum and feces. This proximal-distal gradient in microbial response may be explained by nutrient absorption patterns, as the progressive depletion of dietary lipids, which are primarily absorbed in the proximal small intestine (34). Traditionally, low diversity of the fecal microbiota has been considered a marker of metabolic dysfunction, given its association with high-fat, low-fiber diets and obesity both in mice and humans (35,36). However, recent evidence indicates that, in humans, this pattern does not extend to the proximal parts of the gut. For instance, we previously reported increased microbial richness in the duodenojejunal fluid of individuals with obesity compared to healthy controls, while it was not the case in stool or saliva samples (8). Similarly, other studies have reported higher bacterial counts in the duodenal microbiome of hyperglycemic individuals compared to normoglycemic individuals (37). While diets in humans and in rodent models of diet-induced obesity differ, these results highlight the opposite microbiome dynamics in the small intestine in mice and humans in response to the diet.

In addition, we confirmed the strong bifidogenic effect of fructooligosaccharides. Interestingly, the highest relative abundance of *Bifidobacteriaceae* in FOS-supplemented mice was observed in the ileum (48,7%), supporting the hypothesis that, in mice, fructooligosaccharides are utilized by the microbiome and in particular by bifidobacteria in the small intestine. Consistent with this, a study in rats fed a high-fat, high-sugar diet have shown that FOS supplementation leads to rapid and significant alterations in the small intestinal microbiota, enhancing lipid-sensing mechanisms, food intake regulation and, in fine, energy homeostasis (13). In human, small intestine microbes from ileal fluid obtained from people with an ileostomy have been reported to be able to ferment FOS *in vitro*, achieving 29–89% hydrolysis of FOS with a degree of polymerization between 2 and 10 within five hours (38). Fructooligosaccharides with a low degree of polymerization were metabolized more quickly than those with longer chains. However, most studies relying on the direct analysis of human samples indicate minimal or no fermentation of FOS in the small intestine (39,40), though prebiotic intake significantly decreases small intestinal transit time (41). Although the effect of fructo-oligosaccharides on the composition of the small intestinal microbiome in humans remains to be documented, the interspecies differences in FOS fermentation in the small intestine underscore the translational validity of FOS supplementation effects observed in rodents data to human physiology.

We report that, in mice, the composition of the gut microbiota is more influenced by the dietary composition and prebiotic supplementation than by the ecological niche, in contrast to our primary hypothesis. Diet-induced shifts on the distal microbiome were expected and coherent with the literature as David and colleagues demonstrated that dietary changes could lead to rapid and profound alterations in gut microbiota composition (43). We initially hypothesized that environmental variations, such as pH, oxygen levels, and nutrient availability along the digestive tract, would result in a greater variation in microbiota composition between gut segments. However, unlike humans, in which we showed that only 4% of the bacterial taxa are shared between the duodenojejunal fluid and stool (8), mice appear to exhibit a relatively homogeneous composition of the microbiota throughout the digestive tract. Specifically, 37% of the taxa were detected both in the jejunum and in the feces of the mice. The proportion of shared taxa between the proximal and distal parts of the digestive tract in mice could be underestimated in the present study, as the limited sequencing depth in the jejunum and ileum of the mice did not allow the detection of low abundance taxa. A key factor explaining this homogeneity is likely by the coprophagic behavior of mice, which reintroduces fecal microorganisms in the proximal part of the small intestine. Supporting this hypothesis, previous studies have demonstrated that preventing coprophagy significantly reduces the abundance in the small intestine of anaerobic bacterial groups that we detected in significant proportion in the jejunum and ileum of chow-diet fed mice, such as *Lachnospiraceae*, *Bacteroidales*, and *Erysipelotrichales* (44).

Considering the difficulty of accessing small intestinal samples in humans and the growing interest in small intestinal microbiota research, numerous studies to explore the role of the small intestine microbiome using mice models are to be expected in the future. Our findings underline the differential dynamics of the small intestinal microbiota in human and mice, such as inverse associations between microbial diversity and obesity, distinct FOS fermentation patterns, and greater gut microbiota homogeneity in mice, which pose challenges to translating findings across species. The impact of coprophagy on small intestinal microbiota composition warrants further investigation, as preventing this behavior may improve the translational relevance of rodent models. Additionally, non-coprophagic models could provide a more accurate framework for studying the small intestinal microbiota and its role in metabolic health and disease.

In summary, we show that high-fat diet and FOS supplementation strongly influence gut microbiota composition along the intestinal tract in mice, overriding the anatomical variation that is prominent in humans. Thus, our work underscores the need for careful consideration when extrapolating rodent data to human physiology and highlights the importance of developing models that better mimic human small intestinal microbiota dynamics.

## MATERIAL AND METHODS

### Animal models

23 male C57BL/6J SPF mice (Laboratoires Charles River, France) aged 8 weeks were housed in a controlled environment (temperature 22°C ± 2°C, 12-hour day/night cycle) with free access to food and water. Mice were acclimated for 7 days to a control diet (chow diet: 3.145 kcal/g, 3.1% lipids, 60% carbohydrates. A04. Safe Diets, Germany) without manipulation. After acclimatisation, 3 groups of mice of similar weight and body composition were constituted. Mice in one of the three groups were fed the chow diet (A04-10, Safe Diets) for 15 weeks, while those in the other two groups were fed a lipid-enriched diet (HFD: 5.24 kcal/g, 60% lipids, 20% carbohydrates. D12492, Research Diets, USA). The prebiotic (FOS, Orafti P95, Germany) was administered to the animals via their drinking water (diluted to a concentration of 10% (w/w)). The drinking solution was filtered (0.22 µm filter) and replaced every two days to prevent degradation of the compound. Gloves were changed

The experiments were conducted in strict compliance with the recommendations of Directive 2010/63/EU of the European Parliament and of the Council of September 22, 2010, on the protection of animals used for scientific purposes and were validated by the Comité d’Éthique pour l’Expérimentation Animale n°5 and the French Research Ministry under agreement number #21011.

### Weight and body composition analysis

Mice were weighed once a week and body composition (fat mass, lean mass, and fluids) was determined by nuclear magnetic resonance (Minispec LF50, Bruker, USA).

### Fasting blood glucose and insulin measurements

Animals were fasted for 6 h, then blood glucose was measured from a drop of blood taken from the tail. Plasma insulin concentration was measured using an ELISA kit (Mouse Ultrasensitive Insulin ELISA, ALPCO, USA) from blood samples collected after the 6 hours fast.

### Insulin tolerance test (ITT)

Mice fasted for 6 hours were injected intraperitoneally with human insulin (0.5 UI/Kg body weight). Blood glucose level were measured using a glucometer (Accu Check ROCHE) at times -15, 0, 15, 30, 60 and 90 min after insulin injection.

The HOMA-IR index was calculated using the following formula:

HOMA-IR = (fasting blood glucose (mg/dL) × fasting blood insulin (µU/mL)) / 405.

### DNA extraction from digestive contents and feces

Protocol was adapted from Godon et al.(45) Samples were weighed, then 250 μl of 4M guanidine thiocyanate, 350 μl of 6% sarcosine and 30 μl of DTT were added. Once homogenized, the contents were incubated for 15 minutes at 95°C. To enhance cell lysis, 0.1 and 2 mm diameter silica beads (MP Biomedicals) were added to the samples and samples were then shaken using Precellys 24 Evolution (Bertin Technologies) at a frequency of 10000 hz 6 times for 30 seconds, interspersed with a 30-second pause.

20 mg of PVPP (polyvinylpolypyrrolidone) and 600 μl of TEN-PVPP buffer (a solution containing PVPP, Tris HCl, EDTA and NaCl) were added to the samples. Samples were centrifuged and the supernatant recovered. The samples were washed and the supernatant recovered two additional times. One volume (1:1) of phenol, chloroform, isoamyl alcohol (25:24:1) was added to the supernatants. The tubes were vortexed, centrifuged and the aqueous phase recovered. Samples were then incubated for 1h at 60°C with 15 μl of proteinase K (19 mg/mL, Thermo Fisher), then nucleic acids were precipitated with isopropanol. DNA was then pelleted through centrifugation and resuspended in phosphate buffer and potassium acetate. Samples were then incubated for 45 minutes at 37°C with RNase A (PureLink Thermo Fisher). DNA was then precipitated with absolute ethanol and sodium acetate, pelleted and washed with 70% ethanol three times. The pellets were air-dried and resuspended in 60 μl of TE buffer. For stool samples, DNA was extracted using the same protocol and further purified using PureLink Microbiome DNA columns (Invitrogen) following the supplier’s protocol. DNA was assessed using the Qubit 4 fluorometer (Invitrogen) for quantity and the NanoDrop (Thermofisher) for quantity and quality.

### Sequencing of DNA

Extracted DNA was sequenced using MinION technology from Oxford Nanopore Technologies (ONT) as previously described (24). DNA libraries were prepared using a ligation sequencing kit (SQK-LSK 109, ONT), enabling simultaneous sequencing of samples (12 EXP-NBD104, ONT).

### Bioinformatic data processing (Nanopore)

According to the pipeline we previously developed, the basecalling of the sequencing reads was conducted using the Guppy version 6.0 software (24, https://git.ummisco.fr/ebelda/nanopore.v2.0). To assess the quality of the sequencing run, we utilized files generated during the basecalling process that contain key quality control metrics, including the number of active channels per minute, the yield of the run-in terms of the number of reads, and the read length distributions.

Individual Nanopore reads were taxonomically classified using Centrifuge (version 1.03). For this mouse microbiome analysis, we used the CMMG (Comprehensive Mouse Microbiome Genome) catalog, which contains 1,573 representative prokaryotic genomes derived from the CMMG project.

After classification of the reads by Centrifuge, host-derived sequences were removed. The remaining sequences were then aligned against the corresponding reference genomes from the Centrifuge database using Minimap2 (version 2.24).

The aligned sequences that passed the filtering step were retained for further analysis. We finally constructed an abundance table of species that was then combined with the experimental metadata and a reference taxonomic table reconstructed from Centrifuge’s NCBI taxid identifiers.

### Microbiome data analysis

Downstream analyses were performed using inhouse scripts and Rhea scripts, “a transparent and modular R pipeline for microbial profiling based on 16S rRNA gene amplicons” (46), within an R environment (R version 4.2.2).

We obtained an average of 152,278 reads per sample, with large discrepancies between ecological niches as the average number of sequences per sample was 39,546 in the jejunal content, 108,545 in the ileal content, 189,842 in the cecal content, and 239,869 in the feces **(Fig. S1a)**. One jejunal sample was excluded from subsequent analyses due to very low read count (2,629). Reads were assigned to 1,336 taxa and zero values were imputed to taxa with counts below 3 reads. Their distribution along the gastrointestinal tract was uneven, with an average of 841 taxa for all fecal samples and an average of 335 taxa for all jejunal samples.

To assess whether sequencing depth was sufficient and whether taxonomic coverage was comparable across different ecological niches, rarefaction curves were generated. In most samples, these curves reached an asymptomatic plateau, suggesting that the sequencing depth is sufficient to characterize the microbial ecosystem (**Fig. S1c**). This observation was corroborated by the low values of the slopes at the end of the curve, calculated as the change in species count between the last and the fifth-to-last subsample points and dividing this by the difference in sequencing depth (number of reads) between these points (**Fig. S1b**). However, the differences of slopes at the end of the rarefaction curves in each ecological niches were statistically significant and inversely proportional to the total number of sequences, indicating an uneven taxonomic coverage.

Consequently, α-diversity analyses will be conducted on a “niche-by-niche” basis, rather than across the entire dataset, to avoid biases arising from uneven taxonomic coverage. For β-diversity and relative abundance analyses, bacterial communities can be compared across niches despite uneven sequencing depth, as these methods are based on relative abundances. Prior conducting these analyses, the data were normalized by dividing the number of sequences assigned to each taxon by the total number of sequences in the sample, followed by scaling based on the sample with the lowest sequence count (47,48).

dbRDA (distance-based redundancy analysis) and nMDS (non-metric multidimensional scaling) analysis analyses were performed, with a Bray-Curtis matrix of species as input. Venn diagram analyses were performed using the VennDiagram package after excluding rare taxa, defined as those detected in fewer than 20% of the samples within each mice group.

### Statistical analysis

Statistical analyses were performed using R software. Prior to hypothesis testing, normality of residuals was assessed using the Shapiro-Wilk test, and homogeneity of variances was tested using Bartlett’s test. For variables failing these assumptions, a logarithmic transformation was applied. If assumptions were still not met post-transformation, non-parametric methods were used.

For comparisons involving normally distributed and homoscedastic data across more than two groups, we used one-way or two-way ANOVA as appropriate, followed by Tukey’s Honest Significant Difference (HSD) test for multiple pairwise comparisons. When data violated parametric assumptions, Kruskal–Wallis tests were used, followed by Dunn’s test with Benjamini–Hochberg correction for controlling the false discovery rate (FDR).

In analyses involving repeated measures or grouped data, such as comparisons between different intestinal segments (ecological niches) from the same animal, we employed linear mixed-effects models. We selected this model for its ability to account for the non-independence of samples collected from the same animal, thereby improving the accuracy of statistical estimates and reducing the risk of type I errors related to pseudo-replication. These models included ‘mouse ID’ as a random effect to account for within-subject dependency, and ‘diet’, ‘prebiotic supplementation’, and ‘intestinal segment’ as fixed effects. Where applicable, interaction terms (e.g., diet × segment) were tested. Model residuals were inspected visually (QQ plots, residual vs. fitted) to assess model fit.

For the dbRDA (distance-based Redundancy Analysis), variance partitioning was quantified to evaluate the proportion of total inertia (Bray–Curtis dissimilarity) explained by each phenotypic or experimental variable (e.g., diet, body weight, fat mass). The statistical significance of the explanatory variables in dbRDA was assessed using permutation-based ANOVA (anova.cca function from the vegan R package). Variables with p-values (or FDR-adjusted q-values) ≤ 0.05 were considered statistically significant.

### Comparative study of the duodenal-jejunal and fecal microbiota in humans

Duodenojejunal and fecal microbiota composition of individuals with and without obesity were retrieved from Steinbach *et al.* study^8^. Briefly, duodenal-jejunal fluid (DJF) and stools were obtained participants with obesity and 15 participants without obesity, matched for age and sex. Each participant underwent an endoscopy, during which duodenojejunal fluid was collected, either part of routine care due to mild epigastric pain (in lean subjects) or performed as part of preoperative assessment before surgery (in participants with obesity).

## Data availability

All data generated in this study can be obtained upon request from the corresponding authors. the data for this study have been deposited in the European Nucleotide Archive (ENA) at EMBL-EBI under accession number PRJEB88878.

## ACKNOWLEDGEMENTS AND FUNDING

The authors also thank the Sorbonne Université Animal facilities (UMS28, Paris, France) for their help in carrying out the murine experiments. This study received the following grants: Inserm (International Research Project grant, IRP Parasum), Société Francophone du Diabète (SFD), Leducq foundation, FHU PaCeMM and Fondation pour la Recherche Médicale (FDT201904008276, FDT202106012793). The authors gratefully acknowledge the philanthropic support provided by Quercia Venture Management to KC’s research group, through the Sorbonne University Foundation. Paul Taillandier is supported by a Ph.D. Fellowship from the French Ministère de l’enseignement supérieur, de la recherche et de l’innovation. No funders had any role in the design of the study, data collection, analysis, interpretation, or in manuscript writing.

## COMPETING FINANCIAL INTERESTS

The authors declare that they have no competing interest related to this study.

---

**Figure S1.**
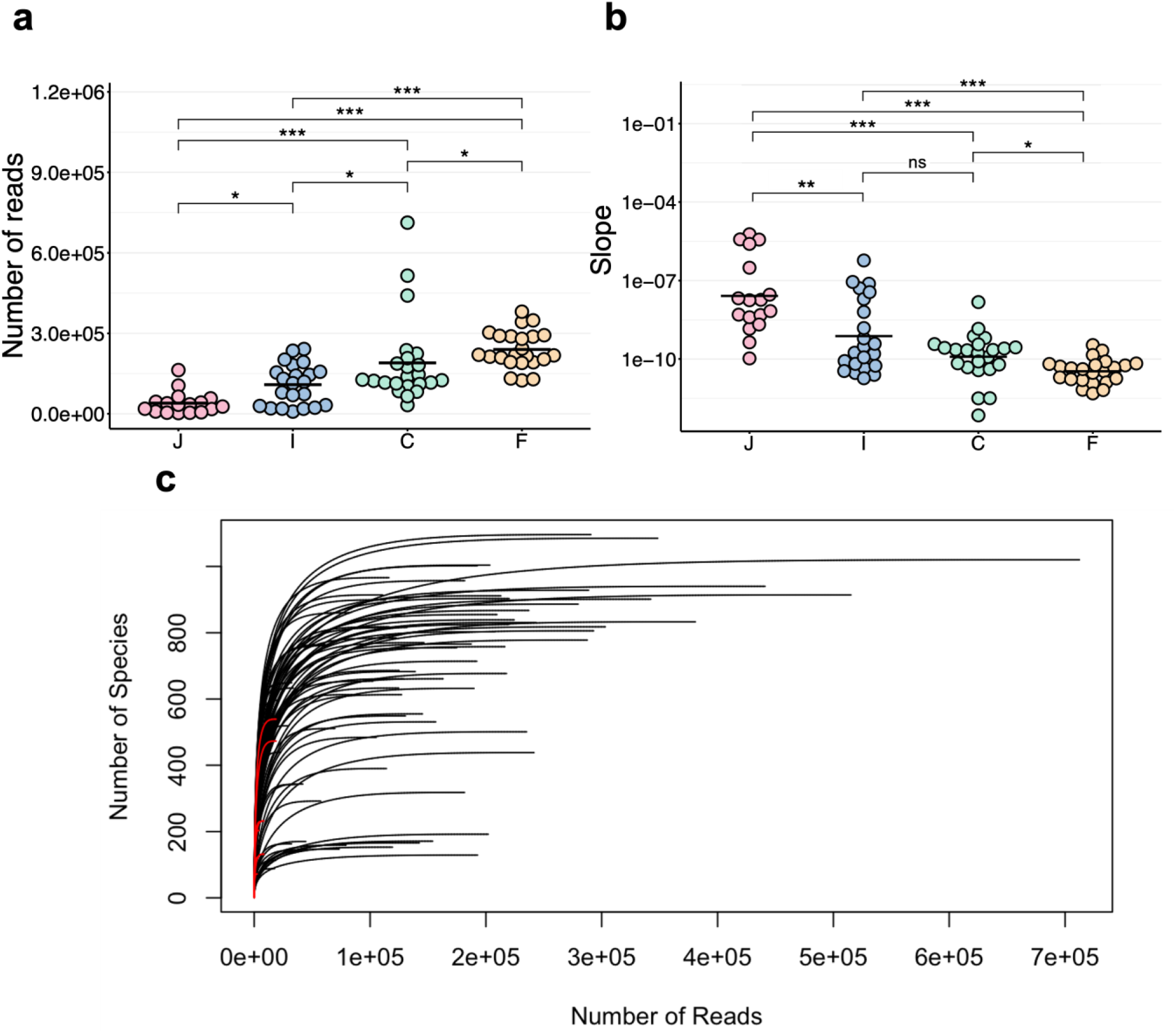
<u>Sequencing depth and taxonomic coverage.</u> (**a**) Average number of reads per sample in the jejunal, ileal, cecal contents, and feces. (**b**) Terminal slopes, calculated as the number of species per 100 sequences, represent taxonomic coverage across niches as illustrated by the rarefaction curves. (**c**) Rarefaction curves for all samples, showing the relationship between sequencing depth and species richness. Data were analyzed using Kruskal–Wallis test followed by Dunn’s pairwise multiple comparison procedure. ns: p > 0.05, *: p <0.05; **: p <0.01; ***: p<0.001.

**Figure S2.**
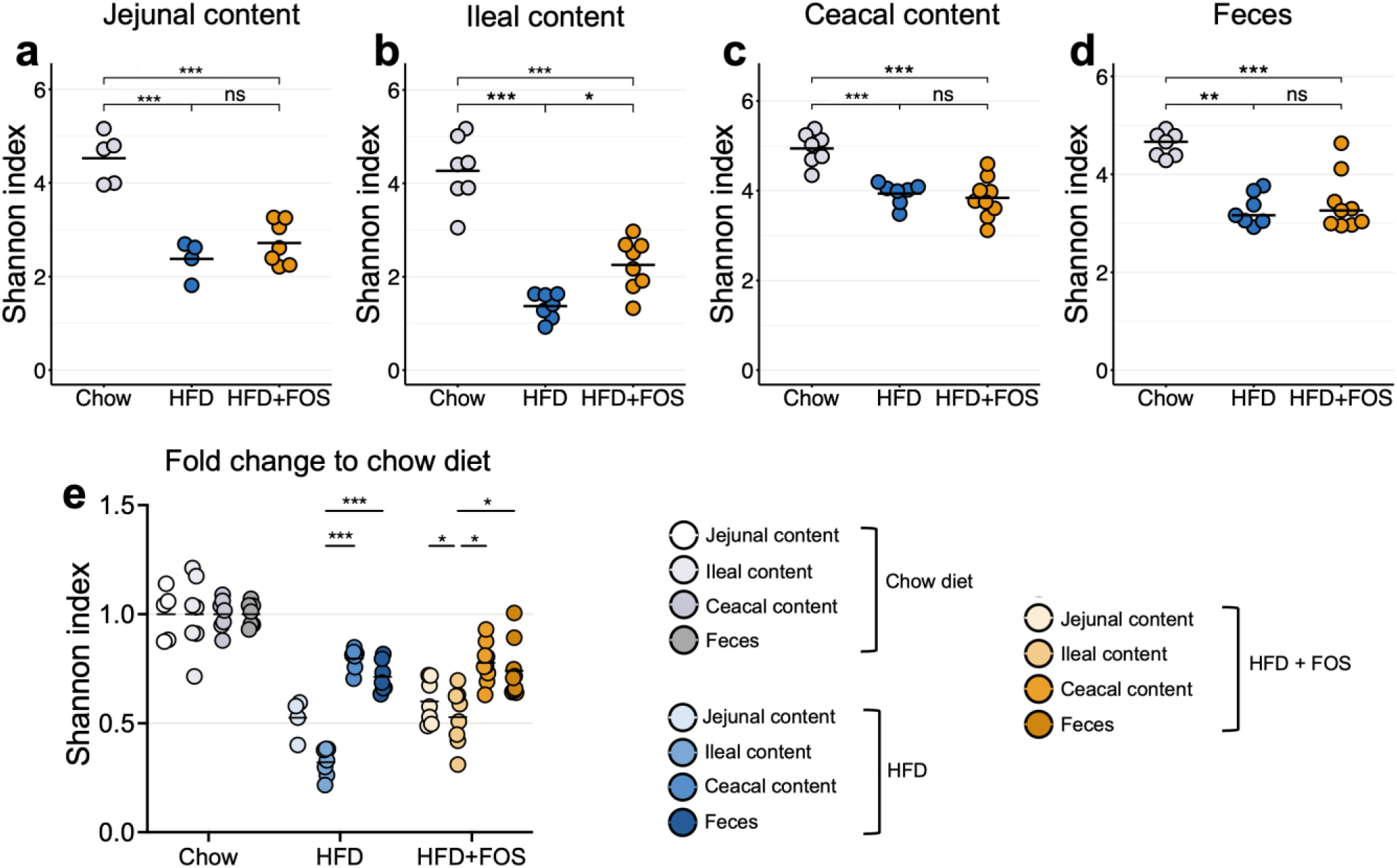
<u>High-fat diet and fructooligosaccharides supplementation modulate the alpha-diversity of the microbiome along the digestive tract.</u> (**a-d**) Shannon index of the jejunal content, ileal content, cecal content and feces of mice fed a chow diet or a HFD and supplemented with fructooligosaccharides (FOS) in drinking water (10% weight/vol) for 13 weeks. The fold change was defined as the ratio between the measured value for (**e**) Shannon index in the HFD or HFD.FOS groups and the mean of the values of the same index observed in the chow diet group. Number of samples per group: 4–9. Results for A, B, C and E are represented as dot plots with means, while the results for the panel D are represented as medians. Data for A, B, C, and E were analyzed using ANOVA followed by Tukey’s post hoc test, while the data for the panel C were analyzed using Kruskal–Wallis test followed by Dunn’s pairwise multiple comparison procedure. ns: p > 0.05, *: p ≤ 0.05; **: p ≤ 0.01; ***: p ≤ 0.001.

**Figure S3.**
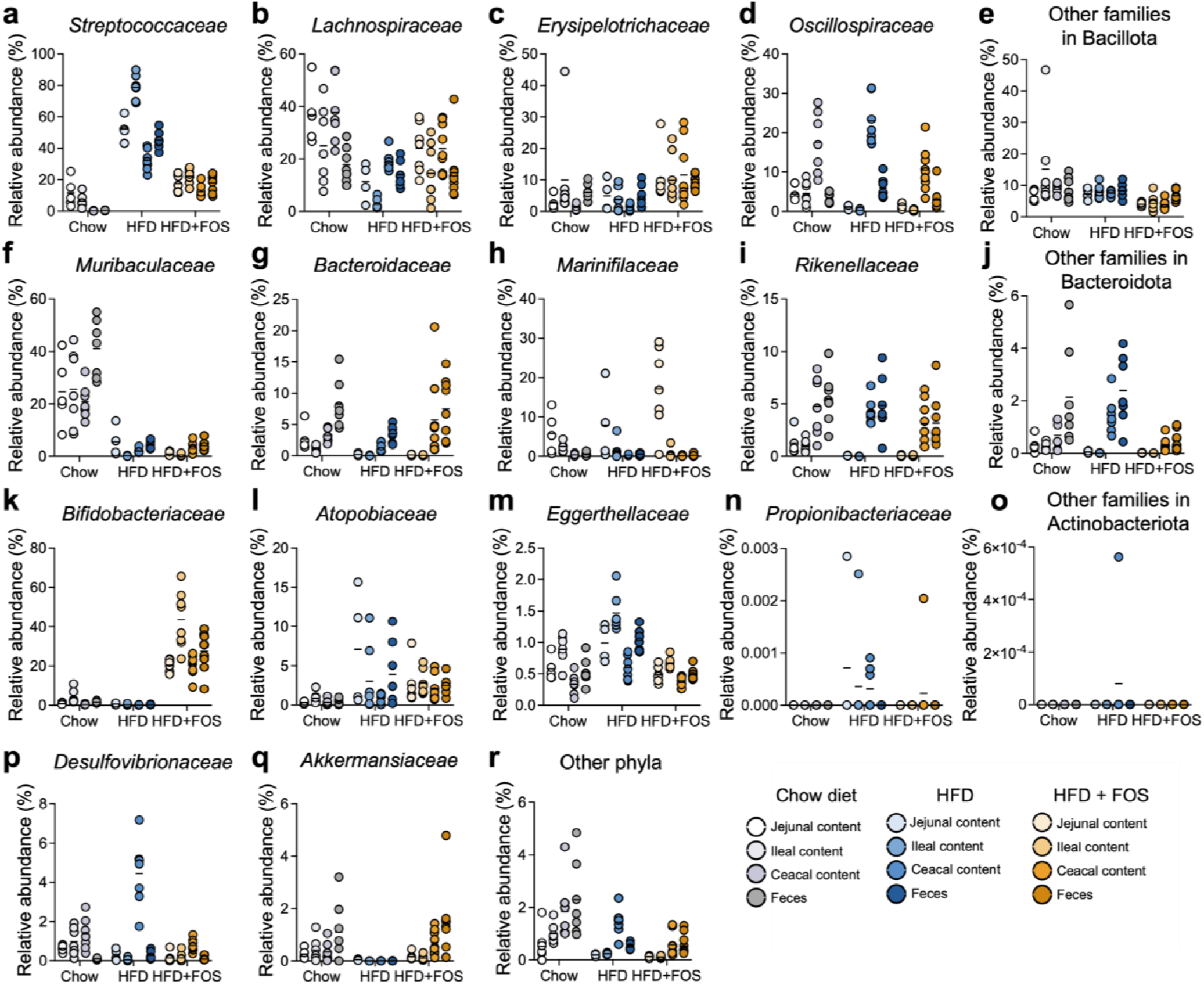
<u>The relative abundance of the most abundant families within the five most represented phyla</u> across all samples was compared between samples from different intestinal segments (Jejunum, Ileum, Caecum, Feces) within the same experimental group, as well as between samples from similar ecological niches but from different groups. Number of samples per group: 4–9. Tukey’s multiple comparisons test was applied after adjusting for a mixed model.

**Figure S4.**
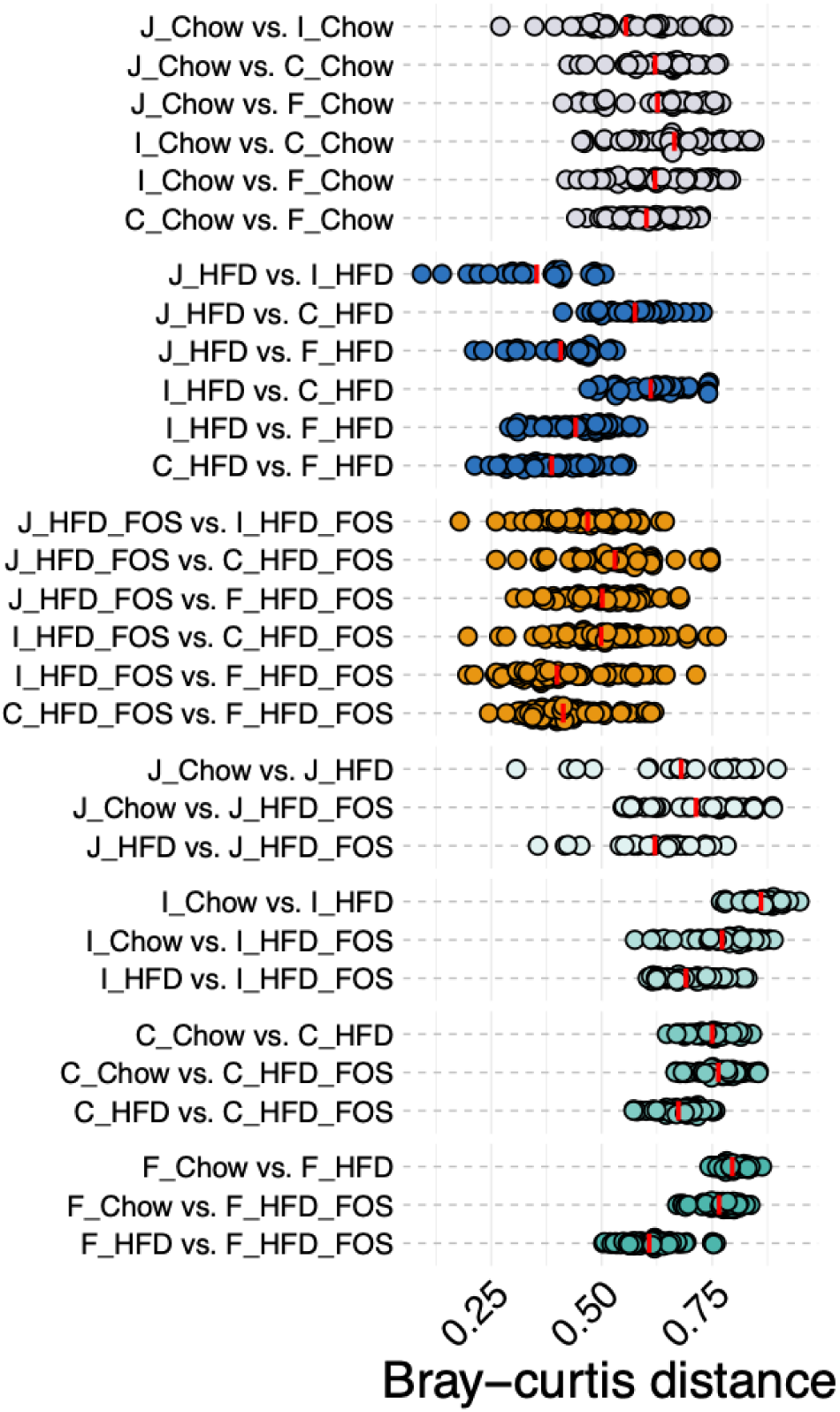
Bray-Curtis dissimilarity distances between pairs of samples from different ecological niche within the same experimental group and between pairs of samples from different experimental groups within each ecological niche. Number of samples per group: 4–9.

## Notes

https://www.ebi.ac.uk/ena/browser/view/PRJEB88878

### Competing Interest Statement

The authors have declared no competing interest.

## REFERENCES

1. Sender R, Fuchs S, Milo R. Revised Estimates for the Number of Human and Bacteria Cells in the Body. PLOS Biol. 19 août 2016;14(8):e1002533.

2. Almeida A, Nayfach S, Boland M, Strozzi F, Beracochea M, Shi ZJ, et al. A unified catalog of 204,938 reference genomes from the human gut microbiome. Nat Biotechnol. janv 2021;39(1):105-14.

3. Jandhyala SM. Role of the normal gut microbiota. World J Gastroenterol. 2015;21(29):8787.

4. Le Chatelier E, Nielsen T, Qin J, Prifti E, Hildebrand F, Falony G, et al. Richness of human gut microbiome correlates with metabolic markers. Nature. 1 août 2013;500(7464):541-6.

5. Manor O, Dai CL, Kornilov SA, Smith B, Price ND, Lovejoy JC, et al. Health and disease markers correlate with gut microbiome composition across thousands of people. Nat Commun. 15 oct 2020;11(1):5206.

6. Aron-Wisnewsky J, Prifti E, Belda E, Ichou F, Kayser BD, Dao MC, et al. Major microbiota dysbiosis in severe obesity: fate after bariatric surgery. Gut. 1 janv 2019;68(1):70.

7. Sundin OH, Mendoza-Ladd A, Zeng M, Diaz-Arévalo D, Morales E, Fagan BM, et al. The human jejunum has an endogenous microbiota that differs from those in the oral cavity and colon. BMC Microbiol. 17 juill 2017;17(1):160.

8. Steinbach E, Belda E, Alili R, Adriouch S, Dauriat CJG, Donatelli G, et al. Comparative analysis of the duodenojejunal microbiome with the oral and fecal microbiomes reveals its stronger association with obesity and nutrition. Gut Microbes. 31 déc 2024;16(1):2405547.

9. Lee JY, Tsolis RM, Bäumler AJ. The microbiome and gut homeostasis. Science. 2022;377(6601):eabp9960.

10. Meier KHU, Trouillon J, Li H, Lang M, Fuhrer T, Zamboni N, et al. Metabolic landscape of the male mouse gut identifies different niches determined by microbial activities. Nat Metab. 1 juin 2023;5(6):968-80.

11. Gibson GR, Hutkins R, Sanders ME, Prescott SL, Reimer RA, Salminen SJ, et al. Expert consensus document: The International Scientific Association for Probiotics and Prebiotics (ISAPP) consensus statement on the definition and scope of prebiotics. Nat Rev Gastroenterol Hepatol. 1 août 2017;14(8):491-502.

12. Dewulf EM, Cani PD, Claus SP, Fuentes S, Puylaert PG, Neyrinck AM, et al. Insight into the prebiotic concept: lessons from an exploratory, double blind intervention study with inulin-type fructans in obese women. Gut. 1 août 2013;62(8):1112.

13. Weninger SN, Herman C, Meyer RK, Beauchemin ET, Kangath A, Lane AI, et al. Oligofructose improves small intestinal lipid-sensing mechanisms via alterations to the small intestinal microbiota. Microbiome. 2 août 2023;11(1):169.

14. Berthoud HR, Kressel M, Raybould HE, Neuhuber WL. Vagal sensors in the rat duodenal mucosa: distribution and structure as revealed by in vivo DiI-tracing. Anat Embryol (Berl). 1 mars 1995;191(3):203-12.

15. Delbaere K, Roegiers I, Bron A, Durif C, Van de Wiele T, Blanquet-Diot S, et al. The small intestine: dining table of host–microbiota meetings. FEMS Microbiol Rev. 1 mai 2023;47(3):fuad022.

16. Le Gall M, Thenet S, Aguanno D, Jarry AC, Genser L, Ribeiro-Parenti L, et al. Intestinal plasticity in response to nutrition and gastrointestinal surgery. Nutr Rev. 1 mars 2019;77(3):129-43.

17. Araújo JR, Tomas J, Brenner C, Sansonetti PJ. Impact of high-fat diet on the intestinal microbiota and small intestinal physiology before and after the onset of obesity. Biochimie. 1 oct 2017;141:97-106.

18. Birchenough GMH, Schroeder BO, Sharba S, Arike L, Recktenwald CV, Puértolas-Balint F, et al. Muc2-dependent microbial colonization of the jejunal mucus layer is diet sensitive and confers local resistance to enteric pathogen infection. Cell Rep. 28 févr 2023;42(2):112084.

19. Tomas J, Mulet C, Saffarian A, Cavin JB, Ducroc R, Regnault B, et al. High-fat diet modifies the PPAR-γ pathway leading to disruption of microbial and physiological ecosystem in murine small intestine. Proc Natl Acad Sci. 4 oct 2016;113(40):E5934-43.

20. Martinez-Guryn K, Hubert N, Frazier K, Urlass S, Musch MW, Ojeda P, et al. Small Intestine Microbiota Regulate Host Digestive and Absorptive Adaptive Responses to Dietary Lipids. Cell Host Microbe. 11 avr 2018;23(4):458–469.e5.

21. Araújo JR, Tazi A, Burlen-Defranoux O, Vichier-Guerre S, Nigro G, Licandro H, et al. Fermentation Products of Commensal Bacteria Alter Enterocyte Lipid Metabolism. Cell Host Microbe. 11 mars 2020;27(3):358–375.e7.

22. Vonaesch P, Araújo JR, Gody JC, Mbecko JR, Sanke H, Andrianonimiadana L, et al. Stunted children display ectopic small intestinal colonization by oral bacteria, which cause lipid malabsorption in experimental models. Proc Natl Acad Sci. 11 oct 2022;119(41):e2209589119.

23. Cani PD, Neyrinck AM, Fava F, Knauf C, Burcelin RG, Tuohy KM, et al. Selective increases of bifidobacteria in gut microflora improve high-fat-diet-induced diabetes in mice through a mechanism associated with endotoxaemia. Diabetologia. 1 nov 2007;50(11):2374-83.

24. Alili R, Belda E, Le P, Wirth T, Zucker JD, Prifti E, et al. Exploring Semi-Quantitative Metagenomic Studies Using Oxford Nanopore Sequencing: A Computational and Experimental Protocol. Genes. oct 2021;12(10):1496.

25. Helander HF, Fändriks L. Surface area of the digestive tract – revisited. Scand J Gastroenterol. 1 juin 2014;49(6):681-9.

26. Constante M, Libertucci J, Galipeau HJ, Szamosi JC, Rueda G, Miranda PM, et al. Biogeographic Variation and Functional Pathways of the Gut Microbiota in Celiac Disease. Gastroenterology. 1 nov 2022;163(5):1351–1363.e15.

27. Chen RY, Kung VL, Das S, Hossain MS, Hibberd MC, Guruge J, et al. Duodenal Microbiota in Stunted Undernourished Children with Enteropathy. N Engl J Med. 22 juill 2020;383(4):321-33.

28. Belda E, Voland L, Tremaroli V, Falony G, Adriouch S, Assmann KE, et al. Impairment of gut microbial biotin metabolism and host biotin status in severe obesity: effect of biotin and prebiotic supplementation on improved metabolism. Gut. 2022;71(12):2463-80.

29. Everard A, Belzer C, Geurts L, Ouwerkerk JP, Druart C, Bindels LB, et al. Cross-talk between Akkermansia muciniphila and intestinal epithelium controls diet-induced obesity. Proc Natl Acad Sci. 28 mai 2013;110(22):9066-71.

30. Woting A, Pfeiffer N, Hanske L, Loh G, Klaus S, Blaut M. Alleviation of high fat diet-induced obesity by oligofructose in gnotobiotic mice is independent of presence of Bifidobacterium longum. Mol Nutr Food Res. 1 nov 2015;59(11):2267-78.

31. Makki K, Brolin H, Petersen N, Henricsson M, Christensen DP, Khan MT, et al. 6α-hydroxylated bile acids mediate TGR5 signalling to improve glucose metabolism upon dietary fiber supplementation in mice. Gut. 1 févr 2023;72(2):314.

32. Moussa FAH, Brownlee IA. Effect of non-digestible oligosaccharides on body weight in overweight and obese adults: A systematic review and meta-analysis of randomised controlled trials. Food Hydrocoll Health. 15 déc 2023;4:100146.

33. Nakamura Y, Natsume M, Yasuda A, Ishizaka M, Kawahata K, Koga J. Fructooligosaccharides suppress high-fat diet-induced fat accumulation in C57BL/6J mice. BioFactors. 2017;43(2):145-51.

34. Zeng X, Xing X, Gupta M, Keber FC, Lopez JG, Lee YCJ, et al. Gut bacterial nutrient preferences quantified in vivo. Cell. 1 sept 2022;185(18):3441–3456.e19.

35. Turnbaugh PJ, Backhed F, Fulton L, Gordon JI. Marked alterations in the distal gut microbiome linked to diet-induced obesity. Cell Host Microbe. 17 avr 2008;3(4):213-23.

36. Cotillard A, Kennedy SP, Kong LC, Prifti E, Pons N, Le Chatelier E, et al. Dietary intervention impact on gut microbial gene richness. Nature. août 2013;500(7464):585-8.

37. Leite G, Barlow GM, Rashid M, Hosseini A, Cohrs D, Parodi G, et al. Characterization of the Small Bowel Microbiome Reveals Different Profiles in Human Subjects Who Are Overweight or Have Obesity. Am J Gastroenterol. juin 2024;119(6):1141-53.

38. van Trijp MPH, Rösch C, An R, Keshtkar S, Logtenberg MJ, Hermes GDA, et al. Fermentation Kinetics of Selected Dietary Fibers by Human Small Intestinal Microbiota Depend on the Type of Fiber and Subject. Mol Nutr Food Res. 1 oct 2020;64(20):2000455.

39. Cherbut C. Inulin and oligofructose in the dietary fibre concept. Br J Nutr. mai 2002;87 Suppl 2:S159–162.

40. van Trijp MPH, Rios-Morales M, Logtenberg MJ, Keshtkar S, Afman LA, Witteman B, et al. Detailed Analysis of Prebiotic Fructo- and Galacto-Oligosaccharides in the Human Small Intestine. J Agric Food Chem. 25 sept 2024;72(38):21152-65.

41. da Silva MVT, Nunes SS, Costa WC, Sanches SMD, Silveira ALM, Ferreira ÁRS, et al. Acute intake of fructooligosaccharide and partially hydrolyzed guar gum on gastrointestinal transit: A randomized crossover clinical trial. Nutrition. 1 oct 2022;102:111737.

42. David LA, Maurice CF, Carmody RN, Gootenberg DB, Button JE, Wolfe BE, et al. Diet rapidly and reproducibly alters the human gut microbiome. Nature. 23 janv 2014;505(7484):559-63.

43. David LA, Materna AC, Friedman J, Campos-Baptista MI, Blackburn MC, Perrotta A, et al. Host lifestyle affects human microbiota on daily timescales. 25 août 2014;15(7):1-15.

44. Bogatyrev SR, Rolando JC, Ismagilov RF. Self-reinoculation with fecal flora changes microbiota density and composition leading to an altered bile-acid profile in the mouse small intestine. Microbiome. 12 févr 2020;8(1):19.

45. Godon JJ, Zumstein E, Dabert P, Habouzit F, Moletta R. Molecular microbial diversity of an anaerobic digestor as determined by small-subunit rDNA sequence analysis. Appl Environ Microbiol. juill 1997;63(7):2802-13.

46. Lagkouvardos I, Fischer S, Kumar N, Clavel T. Rhea: a transparent and modular R pipeline for microbial profiling based on 16S rRNA gene amplicons. PeerJ. 11 janv 2017;5:e2836.

47. Lagkouvardos I, Fischer S, Kumar N, Clavel T. Rhea: a transparent and modular R pipeline for microbial profiling based on 16S rRNA gene amplicons. PeerJ. 11 janv 2017;5:e2836.

48. McMurdie PJ, Holmes S. Waste Not, Want Not: Why Rarefying Microbiome Data Is Inadmissible. PLOS Comput Biol. 3 avr 2014;10(4):e1003531.

